# A Theory for High-Throughput Genetic Interaction Screening

**DOI:** 10.1101/2022.10.05.510977

**Authors:** Madeline E. McCarthy, William B. Dodd, Xiaoming Lu, Nishi D. Patel, Charlotte V. Haskell, Hugo Sanabria, Mark A. Blenner, Marc R. Birtwistle

**Affiliations:** Department of Chemical and Biomolecular Engineering, Clemson University; Department of Physics and Astronomy, Clemson University; Department of Chemical and Biomolecular Engineering, University of Delaware; Department of Bioengineering, Clemson University

## Abstract

Systematic, genome-scale genetic screens have been instrumental for elucidating genotype-phenotype relationships, but approaches for probing genetic interactions have been limited to at most ∼100 pre-selected gene combinations in mammalian cells. Here, we introduce a theory for high-throughput genetic interaction screens. The theory extends our recently developed Multiplexing using Spectral Imaging and Combinatorics (MuSIC) approach to propose ∼10^5^ spectrally unique, genetically-encoded MuSIC barcodes from 18 currently available fluorescent proteins. Simulation studies based on constraints imposed by spectral flow cytometry equipment suggest that genetic interaction screens at the human genome-scale may be possible if MuSIC barcodes can be paired to guide RNAs. While experimental testing of this theory awaits, it offers transformative potential for genetic perturbation technology and knowledge of genetic function. More broadly, the availability of a genome-scale spectral barcode library for non-destructive identification of single-cells could find more widespread applications such as traditional genetic screening and high-dimensional lineage tracing.

## Introduction

Understanding which genes play essential roles in a cellular or organismal process is crucial to our understanding of biology^1^. This can be accomplished by perturbing genes and observing the corresponding phenotype alterations^2^. This process, when applied in parallel to multiple genes one-at-a-time, is known as genetic screening^3,45–8^. Historically, there have been several methods for performing genetic screens, including Zinc finger nucleases (ZFNs) and transcription activator-like effector nucleases (TALENs) which are engineered nucleases that induce DNA DSBs at specific locations^9,10^, RNAi which uses double stranded RNAs (or a short hairpin (sh)RNA) to knock down the gene-of-interest^11^, and CRISPR which induces DNA breaks or alters transcription at specific sites in the genome^12,13 15^.

While these gene perturbation technologies have revolutionized biomedical science, most genome-scale screens (outside of organisms like *S. cerevisae*^16^) remain limited to one gene at a time^17^. However, often genes cooperate with one another to influence phenotype. Such cooperation is called genetic interaction^18–21^. Recent approaches have made progress towards larger scale genetic interaction screening. For example, cloning two different CRISPR gRNAs into a single plasmid enables interaction screening for ∼100 pre-selected genes ^19,25–27^. Other approaches include dual recombinase-mediated cassette exchange to create mosaic *in vivo* models harboring multiple desired cancer driver mutations^28^, or using protein epitope combinatorial barcodes (pro-codes) with mass cytometry to perform high-dimensional CRISPR screens on 100s of selected genes in single cells^29^. The sheer number of observations that must be made to cover human gene interactions space almost necessitates a single-cell approach, like Perturb-seq^30–32^. However, genetic interaction screening approaches that scale past ∼100 genes have yet to be described.

Here, we propose that our recently developed fluorescence **mu**ltiplexing with **s**pectral **i**maging and **c**ombinatorics (MuSIC)^33^ approach may be compatible with single cell genetic interaction screening that could scale to the full human genome. MuSIC uses combinations of fluorophores (proteins or small molecules) to create spectrally unique MuSIC probes. Here we introduce the concept of further combining MuSIC probes into MuSIC barcodes for increased diversity and thus multiplexing. Moreover, because these spectral barcodes are fluorescence-based, they can be read non-destructively. Theory and simulations based on currently available fluorescent proteins suggests that given a palette of 18 fluorescent proteins, ∼400,000 MuSIC barcodes could be generated, far surpassing human genome-scale. Simulations suggest that given current spectral flow cytometry equipment and experimental noise, human genome-scale genetic interaction screens may be possible. More advanced instrument hardware such as more excitation lasers and/or higher resolution emission spectra could increase such capabilities. While experimental testing of this theory awaits, it offers transformative potential for genetic perturbation technology and knowledge of genetic function. More broadly, the availability of a genome-scale spectral library for non-destructive cell identification could find more widespread applications such as traditional genetic screens and high-dimensional lineage tracing.

## Methods

### Availability, Code Overview, and Simulation

All MATLAB code and raw data used for simulations are included in the supplementary code zip file associated with this manuscript. The scripts `GenerateProbeData_3l_HN.m, GenerateProbeData_3l_LN.m, GenerateProbeData_5l_HN.m,` and `GenerateProbeData_5l_LN.m` are used for generating the list of good probes for single probes, barcodes, and two barcodes, for 3 lasers/high noise (HN), 3 lasers/low noise (LN), 5 lasers/high noise, and 5 lasers/low noise respectively. The core of these scripts is done by the functions `RemoveProbes_onebyone.m, RemoveBarcodes_onebyone.m`, and `RemoveTwoBarcodes_onebyone.m,` respectively. The README file contains relevant information on the code for execution and reproducing the results. These simulations were performed in MATLAB using 40 CPUs on the Palmetto supercomputing cluster at Clemson University.

### Data Sources

Emission spectra, excitation spectra, and brightness for fluorescent proteins were gathered from *fpbase*.*org* (**Supplementary Table 1** and references therein). Specifications for flow cytometer noise, excitation channels, and emission binning were obtained from the Aurora and Northern Lights flow cytometer user guides on *cytekbio.com*.

### Simulated FRET Efficiency and MuSIC Probe Selection

FRET efficiency *ε* between two fluorophores is typically calculated as follows

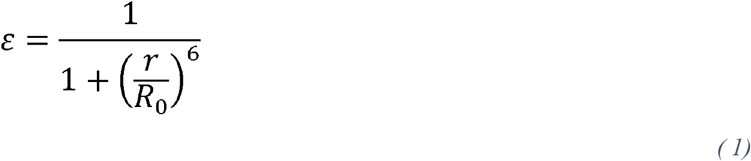

where *r* is the distance between the two fluorophores and *R*_*0*_ is the Förster radius^34^. The Förster radius is the distance between fluorophores that gives a 50% FRET efficiency^34^. Thus, to estimate the FRET efficiency between any given pair of fluorescent proteins, we must calculate *R*_*0*_ and *r*.

The Förster radius can be cast as follows^35^

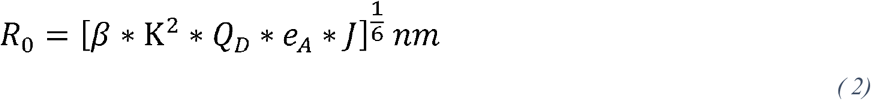

where *β* is a constant (which also converts to nm), K^2^ is an orientation factor between the two fluorophores, *Q*_*D*_ is the donor quantum yield, *e*_*A*_ is the maximal acceptor extinction coefficient (M^-1*^cm^-1^), and *J* is the spectral overlap integral. The value of K^2^ is not usually known (nor easily measurable) but is assumed to be a constant value of 2/3 for isotropic reorientation of the coupled fluorophores ^36^. This value may not be 2/3 for fluorescent protein tandems but in practice, deviations can be accounted for by the constant *β*^37^. J is calculated as follows

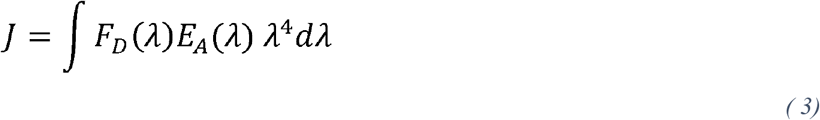

wher*e F*_*D*_ is the normalized emission spectra of the donor and *E*_*A*_ is the normalized excitation spectra of the acceptor, which both are evaluated at wavelength *λ*. Here, the spectral data is normalized to have a maximum value of 1. We calculate the overlap integral using the function `trapz` in MATLAB (see code) with bounds from *λ* = 300 to 800 nm. The value for *β* is estimated to be 6.33*10^−6^ based on a known Förster radius of 6.1 nm for mTFP-Venus^38^ (along with known *Q*_*D*_, and *e*_*A*_, and *J* calculated as above).

The closest physical distance that chromophores of fluorescent proteins can be is ∼3 nm^39^. Furthermore, most high FRET producing pairs have an *R*_*0*_ greater than 5 nm^40^. Thus, we do not consider MuSIC probes that have *R*_*0*_ < 5 nm. Since the distance between fluorescent proteins can usually be adjusted (by linker length, for example), we set *r = R*_*0*_ in simulations, giving a FRET efficiency of 50% for each MuSIC probe with more than one fluorescent protein.

### Simulating Reference Emission Spectra for MuSIC probes

There are three classes of MuSIC probes that require separate consideration for simulating their emission spectra: those made of a (i) single fluorescent protein, (ii) two fluorescent proteins, and (iii) three fluorescent proteins. The below equations are used to generate columns of the reference matrix **R** (see below) for unmixing. Each simulated spectra for a single excitation channel has a value every nm from 300 to 800 nm. The below model assumes that tandem fluorescent proteins have the same properties as the monomers, that static quenching is not a dominant feature, and that fluorescent protein maturation is not a significant factor for the spectra. We assume cross-talk is negligible, but for all intents and purposes, it would be observed as effective FRET-related activity and therefore is expected to not have additional functional consequences for simulation results. We also note here that this model does not take into account detector quantum efficiency. Avalanche Photodiode (APD) detectors (used in the Cytek instruments) generally have slightly lower quantum efficiency in the lower wavelengths (UV/Blue), but so long as all reference spectra and samples are measured with the same instrument, this would not introduce any further bias and not affect the conclusions drawn here.

To simulate the emission intensity spectra *I* for a single fluorescent protein MuSIC probe, given a particular excitation wavelength (*λ*_*ex*_)and vector of emission wavelengths from 300 to 800nm at every nm (*λ*), the following equation is used (adapted from Schwartz et. al)^41^

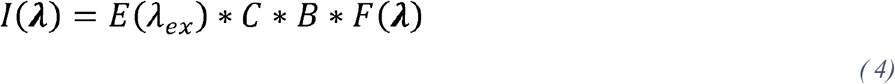

Where *E* is the fraction of excited fluorophores and is a function of excitation wavelength (explained below), *C* is the relative probe concentration (taken as 1 for reference spectra assuming a null condition of equal expression levels between probes), *B* is the brightness (product of maximal extinction coefficient and quantum yield), and *F* is the normalized emission spectra vector of the fluorescent protein (normalized as above). *E(λ*_*ex*_) is given by the fluorescent protein’s normalized excitation spectra at the designated excitation wavelength.

For MuSIC probes with two fluorescent proteins, called 1 and 2 ordered from blue to red, the emission intensity spectra *I*(*λ*) has three contributing components: acceptor emission due to FRET (*I*_*2,1*_), donor emission (*I*_*1*_), and acceptor emission due to direct excitation (*I*_*2*_). The overall emission intensity spectra *I* is the sum of the three components

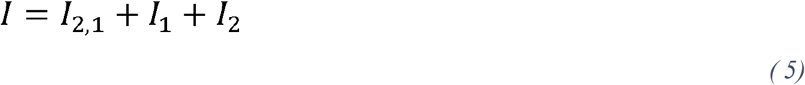

Each of these terms depends on the FRET efficiency. We assume that FRET efficiency is reduced due to any direct acceptor (2) excitation, since excited acceptors would not be able to undergo FRET. This adjusted FRET efficiency, *ε*_*adj*_, is calculated as follows

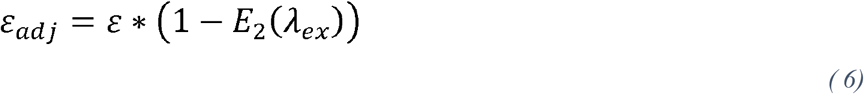

where *E*_*2*_ is the fraction of excited fluorophores for fluorescent protein 2 and the term (1-*E*_*2*_) denotes the fraction of fluorescent protein 2 molecules that have not been directly excited.

Fluorescent protein 2 emission due to FRET from fluorescent protein 1 is then calculated by

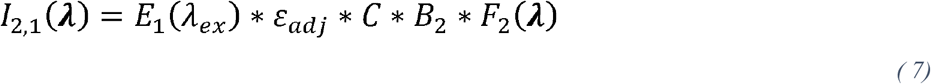

This emission intensity is proportional to emission properties of fluorescent protein 2 (emission spectra and brightness), the fraction of excited molecules for fluorescent protein 1, and the adjusted FRET efficiency between the two fluorescent proteins. Fluorescent protein 1 emission is calculated by

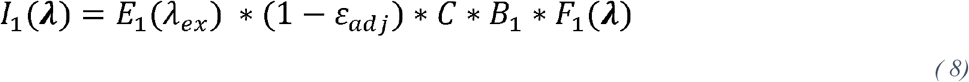

This emission is calculated similarly to that above for a single fluorescent protein; however, it is corrected to only take into account the fraction of excited molecules that are not undergoing FRET (1 − *ε*_*adj*_).

Fluorescent protein 2 emission due to direct excitation is calculated by

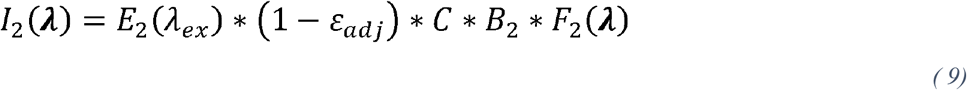

We opt here to be conservative and reduce the amount of fluorescence from direct excitation of fluorescent protein 2 by the FRET taking place.

For MuSIC probes with three fluorescent proteins, called 1, 2, and 3 ordered from blue to red, the emission intensity depends on six different components. Three are due to direct excitation: emission intensity of fluorescent protein 1 (*I*_*1*_), emission intensity of fluorescent protein 2 (*I*_*2*_), and emission intensity of fluorescent protein 3 (*I*_*3*_). The other three are due to FRET: FRET sensitized emission intensity of fluorescent protein 2 due to FRET with fluorescent protein 1 (*I*_*2,1*_), FRET sensitized emission intensity of fluorescent protein 3 due to FRET with fluorescent protein 2 that ultimately came from FRET with fluorescent protein 1 (*I*_*3,1*_), and FRET sensitized emission intensity of fluorescent protein 3 due to FRET with fluorescent protein 2 (*I*_*3,2*_). The overall intensity is calculated as the sum of the six intensities. We assume negligible direct FRET from fluorescent protein 1 to 3.

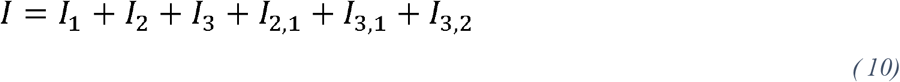

The adjusted FRET efficiencies between fluorescent proteins, *ε*_*adj1*_ and *ε*_*adj2*_, are calculated as above

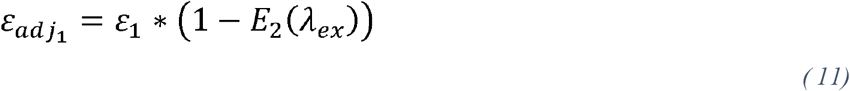

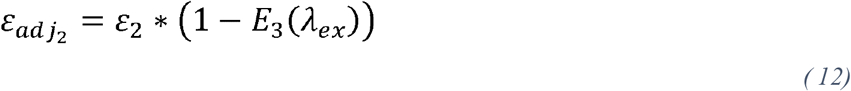

The emission intensity of fluorescent protein 1 due to direct excitation is calculated by

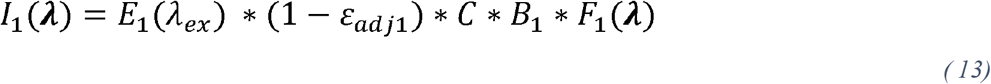

This emission is calculated similarly to that above and is corrected to only consider the fraction of excited fluorescent protein 1 molecules that are not undergoing FRET with fluorescent protein 2.

The emission intensity of fluorescent protein 2 due to direct excitation is calculated by

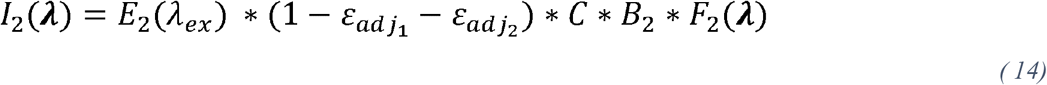

This emission is corrected to only consider the fraction of excited fluorescent protein 2 molecules that are not undergoing FRET with either fluorescent proteins 1 or 3.

The emission intensity of fluorescent protein 3 due to direct excitation is calculated by

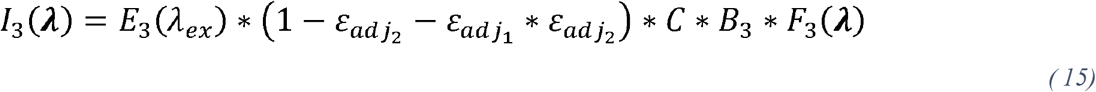

This emission intensity only considers the fraction of fluorescent protein 3 molecules that are not involved in FRET with either fluorescent protein 2 or FRET from the first fluorescent protein through the second.

The emission intensity of fluorescent protein 2 due to FRET from fluorescent protein 1 is calculated by

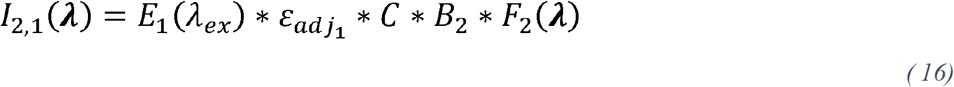

The emission intensity of fluorescent protein 3 due to FRET from fluorescent protein 1 through fluorescent protein 2 is calculated by

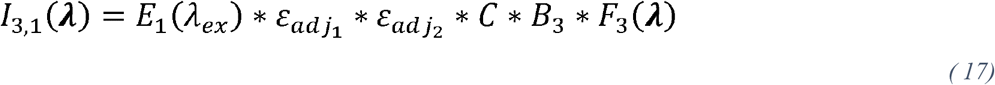

Finally, the emission intensity of fluorescent protein 3 due to FRET from fluorescent protein 2 is calculated as follows

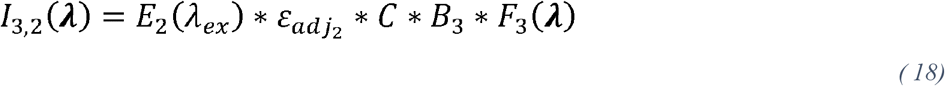

### Calculating the Observed Spectra Using Cytek Binning

The emission spectra of the MuSIC probes are simulated at every nm as described above. To best replicate the emission spectra generated from the Cytek Northern Lights and the Cytek Aurora flow cytometers, we condensed the simulated emission spectra based on the emission channels for each instrument, referred to as binning. Each emission channel represents spectral data condensed over a range of wavelengths, so to convert the simulated emission spectra (which is at every nm) we averaged the simulated emission spectra *I* for each probe over the wavelength ranges of each instrument’s emission channels. Each binned emission point is calculated as follows

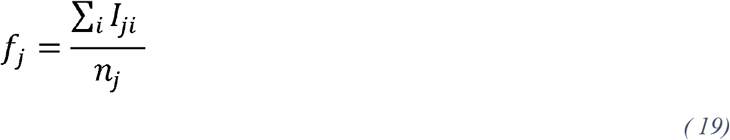

Where *f*_*j*_ is the binned emission point over the wavelength range for channel *j, n* is the number of wavelengths in channel *j*, and *I* is calculated as above.

### Noise Model

Noise is assumed to be normally distributed and simulated using the MATLAB function `randn`. The standard deviation for the normal distribution is estimated based on data from the Cytek Northern Lights flow cytometer, given by the manufacturer, which is estimated at 50 relative fluorescent units (RFUs) for an intensity of 10^5^ RFUs. In the above simulations, the fluorescence emission spectra have an average maximum of ∼10 RFUs. The standard deviation of 50 is thus decreased by a factor of 10^4^ to adjust for the simulated emission spectra, giving a standard deviation of 0.005 for noise. This value is used as the value for “low” noise. The standard deviation is set to 0.05 for “high” noise (10-fold higher than the low noise).

### Unmixing

The fluorescence emission spectra of a mixture of fluorophores can be cast as a sum of the emission spectra of the individual fluorophores as follows.

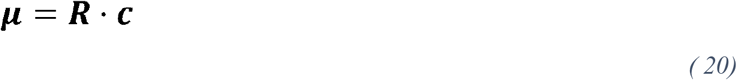

Where ***µ*** is an *n-by-1* vector of observed fluorescence emission intensity at *n* emission wavelength/excitation channel combinations, **R** is an *n-by-m* reference matrix that is generated from the simulated emission spectra of *m* individual probes with multiple excitation channels as described above, and ***c*** is an *m-by-1* vector containing the relative probe concentrations.

Solving this equation gives an estimate of the relative probe concentrations, ***c***. This is done using the MATLAB function `lsqlin`. The lower bound for elements of ***c*** is set to zero, and the upper bound is left empty.

### Generating a Simulated Experimental Data Set

Simulated data are generated by first specifying the relative probe concentrations for different mixtures of MuSIC probes. This is referred to as the actual mixture composition vector, **c**_**a**_. For single probe mixtures, one probe concentration is set to 1 and all others are zero. For barcode mixtures, two probe concentrations are set to 1, and all others are zero. For two barcode mixtures, four probe concentrations are set to 1, and all others are zero. For the case of variable probe expression levels, probe concentrations are set to a random number between 0.5-1.5 (`rand`). For two barcode mixtures, the probes are divided into two batches, and two probes are chosen from the first batch while two are chosen from the second (see Results). Equation 20 with **c**_**a**_ and **R** is used to calculate ***µ***_***a***_, the simulated emission spectra of the mixture. Experimental noise is then added to the simulated emission spectra at either low or high levels, as described above, giving ***µ***_***n***_, the simulated observed spectra. Finally, Equation 20 is used to solve for **c** (i.e., unmixing), giving the predicted mixture composition, **ĉ**.

### Binary Classification

Binary classification is performed on the predicted mixture composition vector by converting the relative level for each probe to a one or zero based on a threshold for each probe. The threshold for each probe is determined as that which gives the maximum Matthews Correlation Coefficient value for each probe respectively based on simulation data (see below).

### Confusion Matrix and Matthews Correlation Coefficient (MCC)

Evaluating binary classification performance requires the calculation of a confusion matrix, which serves as a centralized table that tracks the number of true and false positive and negative classifications. The confusion matrix allows for the calculation of a multitude of performance metrics and is calculated using the MATLAB function `confusionmat.m`. Out of these different metrics, the Matthew’s Correlation Coefficient (MCC), or phi coefficient, was chosen to quantify the performance of probes in the simulations. The MCC was chosen because it is appropriate when the classes are highly imbalanced^42^, such as what we have here when there are many more true negatives than true positives. Other metrics, such as the F1 score or Accuracy, are problematic for situations where there might be significantly more true negatives than false positives.

Given a classification threshold to evaluate, a confusion matrix is generated for each probe using the actual mixture compositions and the binary predicted probe concentrations for each probe. These confusion matrices are used to generate an individual MCC score for each probe, given the threshold. The threshold is then varied to determine the optimum threshold to maximize MCC for a particular probe.

A confusion matrix is generated for the entire group of probes using a matrix of all concatenated actual mixture composition vectors and a matrix of all concatenated predicted mixture composition vectors. This confusion matrix is used to generate the overall MCC score which represents the performance for the entire group of probes.

## Results

This paper explores a theory for creating a large library of genetically-encoded, fluorescence spectral barcodes for potential application to genetic interaction screening. It is based on our recently published Multiplexing using Spectral Imaging and Combinatorics (MuSIC) approach^33^, which creates unique spectral signatures from stably-linked combinations of individual fluorophores. The individual fluorophores or combinations are called MuSIC probes. In this work, we consider expanding the number of fluorophores used by fusing 2 or 3 individual fluorophores that would give rise to unique spectral properties. The spectral signatures of combination probes are linearly independent (i.e., unique) from the individual fluorophore spectra comprising the combination so long as sufficient Förster resonance energy transfer (FRET) occurs. This linear independence property allows for the estimation of individual MuSIC probe levels when they are together in a mixture, a process often called “unmixing”.

We selected 18 fluorescent proteins (see Methods and **Table S1**) that span the ultraviolet to infrared spectrum and first wanted to determine how many MuSIC probes could be generated. The quality of unmixing depends on the FRET efficiency, which is directly related to the Förster radius and the physical distance between chromophores of the fluorescent proteins (see Methods). The distance between fluorescent proteins can usually be adjusted by altering the length and nature of the peptide fusion linker; thus, the answer to this question depends on the Förster radius chosen as acceptable (**Fig. 1A-B**). Since high FRET producing pairs usually have a Förster radius greater than 5 nm^40^, we only consider MuSIC probes that have an estimated Förster radius greater than 5 nm. At this cutoff, 910 MuSIC probes can be generated (**Table S2**), but this is far from genome-scale. We should also note here that in principle the same fluorescent proteins in a probe could be engineered to be a different distance apart and thus a different FRET efficiency, which would increase the number of probes. However, for the purposes of this work, we only consider one FRET efficiency (∼50%) per probe.

**Figure 1:**
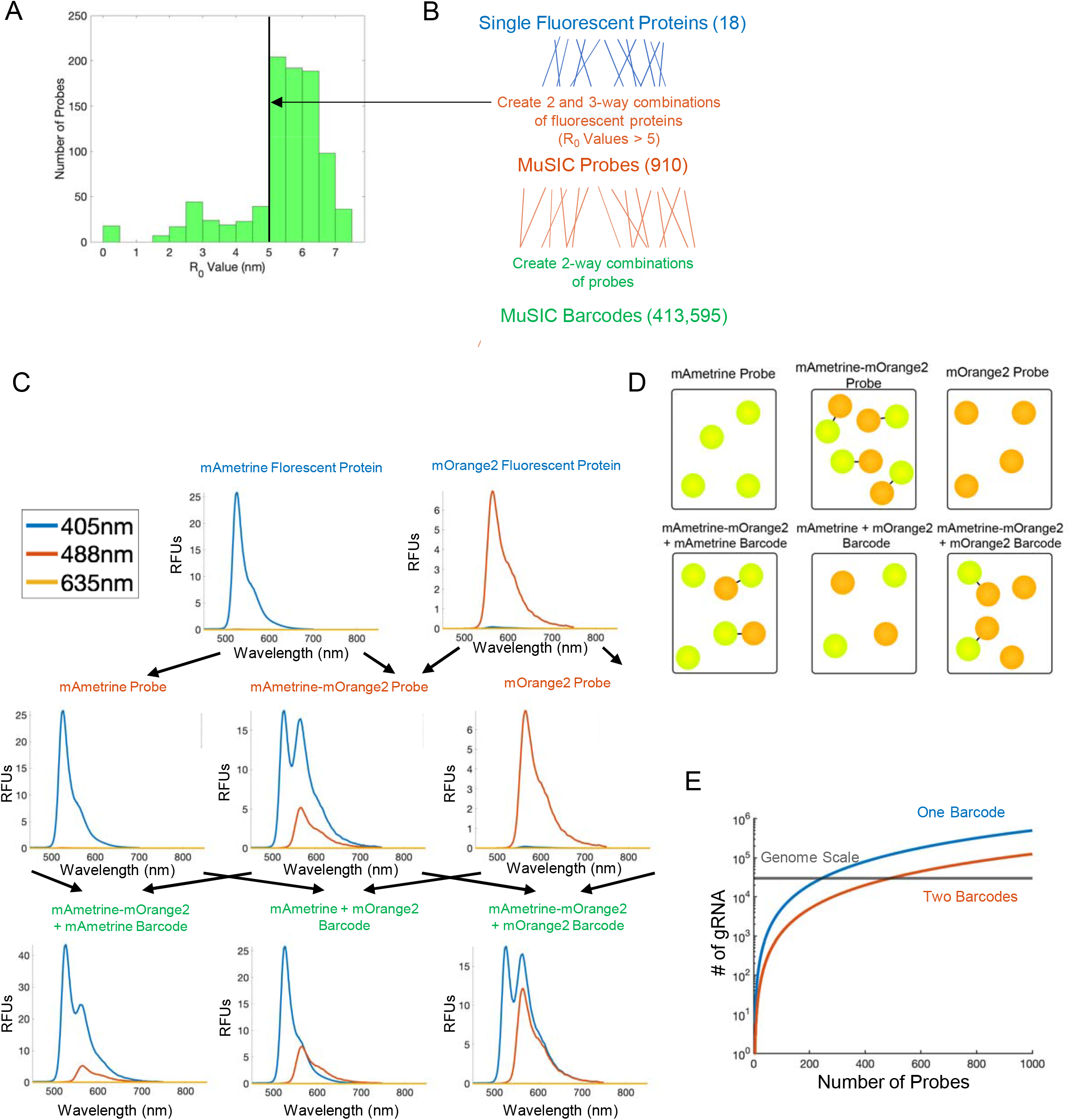
Theory and scope of MuSIC Barcodes for genetic and genetic interaction screening. (A) Forster Radius (R_0_) cut off for probe selection. From the total list of possible MuSIC probes (987), only probes with an R_0_ value greater than 5nm (910) are selected as potentially good probes. (B) Potential number of MuSIC probes and barcodes. Given 18 fluorescent proteins, 910 MuSIC probes can be created (with an R_0_>5nm), and given 910 MuSIC probes, 413,595 MuSIC barcodes could be created. (C) Example emission spectra of MuSIC probes and barcodes when excited at 405, 488, and 635nm. Given the fluorescent proteins mAmetrine and mOrange2, three MuSIC probes can be created that are spectrally unique. Given these three MuSIC probes, three MuSIC barcodes can be created. (D) Schematic showing the creation of MuSIC probes and barcodes from single fluorescent proteins. (E) Genetic and genetic interaction screening capabilities given the number of MuSIC probes that can be created.

Can we develop another layer of combinatorics to generate further diversity? Consider the concept of a MuSIC barcode that is a combination of MuSIC probes. As an example, let us start with two fluorescent proteins, mAmetrine and mOrange2. From these two fluorescent proteins we can create three MuSIC probes: a single fluorescent protein probe of mAmetrine, the combination probe of mAmetrine and mOrange2, and another single fluorescent protein probe of mOrange2. A MuSIC barcode is then every 2-way combination of the probes. Thus, from these probes we can create three MuSIC barcodes **(Fig. 1C-D)**. The MuSIC barcode spectra are clearly unique from one another. The number of barcodes that can be generated given a particular number of probes is given by combinatorics (see Methods); 910 probes gives 413,595 barcodes (**Fig. 1B, E**).

This barcode diversity far exceeds the number of genes in the human genome (**Fig. 1E**). If each MuSIC barcode could be paired to a guide RNA (gRNA), and if resolvable in practice, one could perform genome-scale genetic screening that is non-destructive in single cells. Specifically, then if a certain MuSIC barcode is detected in a particular cell (via a fluorescence emission spectra measurement), that would indicate the gRNA that was present, and therefore the target gene that was likely modulated in that cell.

MuSIC barcodes may also enable large-scale genetic interaction screening (**Fig. 1E**). Consider that a gRNA is paired to a MuSIC barcode as above, but instead there are two MuSIC barcodes in a cell corresponding to two specific gRNAs. This means four MuSIC probes would be present in the cell. To avoid mapping ambiguity from probes to barcodes to gRNA, the MuSIC probe library would have to be split in half before linking gRNA with MuSIC barcodes, which makes the predicted scale of genetic interaction screening lower than that of genetic screens. With 910 MuSIC probes, 103,285 gRNA could be studied for genetic interactions, which approaches human genome-scale genetic interaction screening at ∼3x redundancy.

While the above suggests MuSIC barcodes may enable novel genetic screening technology, how well might it work in practice? Of the 910 potential probes, how many can reliably be identified from expected mixtures? To constrain the answer to this question, we developed a simulation workflow. Rapid measurement of fluorescence emission spectra in single cells has recently become possible with Cytek flow cytometers. For this reason, we have based the simulation studies described in this paper off the Cytek Northern Lights Flow Cytometer (3 lasers; 405, 488, and 635nm) and the Cytek Aurora Flow Cytometer (5 lasers; 356, 405, 488, 561, and 635nm) **(Fig. 2A)**. The spectral emission bin structure for each instrument and its signal-to-noise ratio is known and we incorporate such information into our simulated measurements (**Fig. 2A—**see also Methods). For genetic screens, it can be useful to reserve one excitation channel to measure an observed phenotype. Therefore, we also investigated a setup for 2 lasers (Northern Lights, dedicating the 635nm laser to a phenotype) and 4 lasers (Aurora, dedicating 635nm laser to phenotype) **(Fig. 2B**).

**Figure 2:**
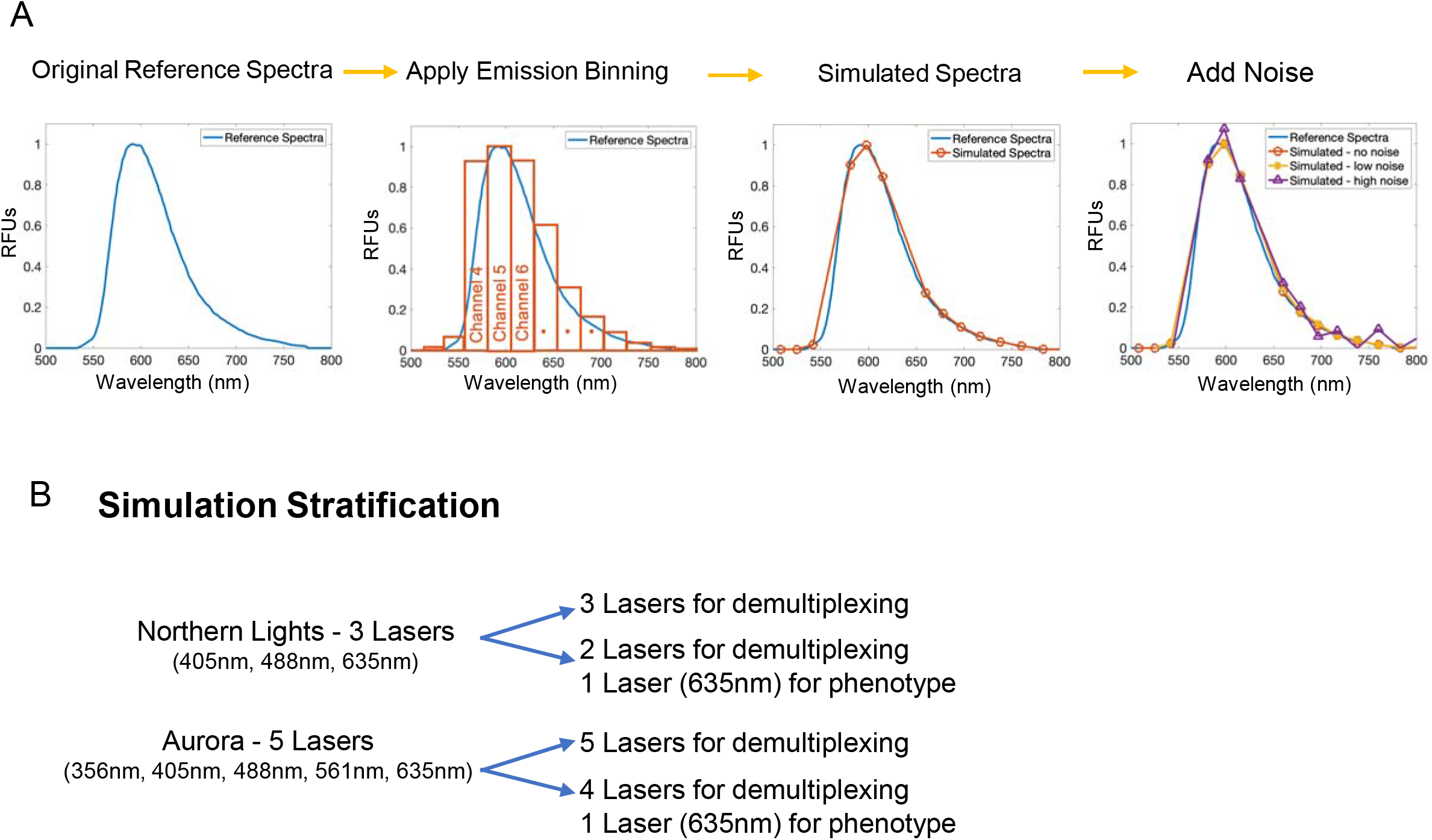
Simulation setup. (A) Simulating emission spectra. Process of condensing the original emission spectra at every nm according to the emission binning and noise of the simulated instrument. (B) Cases for the simulation experiment setup based on Cytek flow cytometers.

We implemented the following simulation strategy to eliminate “poorly” performing probes from consideration **(Fig. 3A)**. A “poorly” performing probe is one that leads to at least one misclassification event in simulations. At the core of the algorithm is a simulated MuSIC probe mixture. This is a vector that represents which probe or probe(s) are present in the ground truth, which we call the actual mixture composition. Using the actual mixture composition vector and the calculated reference matrix (see Methods—spectra of individual probes), we can calculate the emission spectra of the mixture. We add low or high noise (see Methods—based on Cytek flow cytometer specs) to the emission spectra of the mixture, generating the simulated observed spectra. After noise is added, we perform linear unmixing, which generates the predicted mixture composition. To compare the predicted mixture composition to the actual mixture composition, we first perform binary classification (see Methods). To quantify performance, we calculate the Matthews correlation coefficient, which is suitable for cases such as this where there are many more true negatives than true positives. If overall classification is not perfect (MCC < 1), then we identify which probe has the worst MCC, and remove it. The simulation is repeated until overall classification is perfect **(Fig. 3B)**, at which point we obtain the final list of good probes (**Table S2**). This process is performed in triplicate.

**Figure 3:**
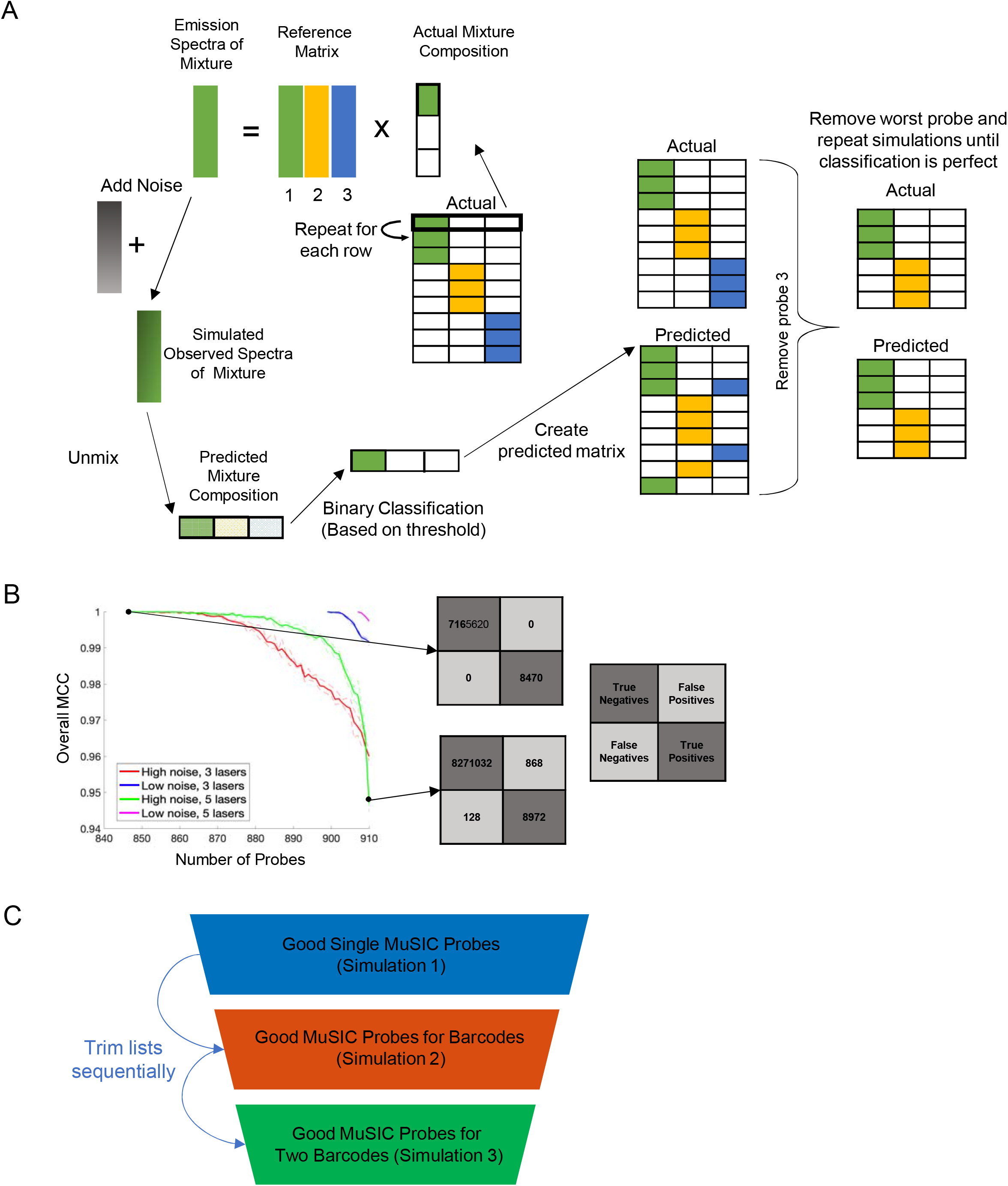
Workflow for probe removal. (A) Obtaining the list of good probes based on classification metrics. First, the emission spectra of a mixture of probes is simulated given a set of probes. Next, noise is added to the emission spectra and the spectra is unmixed (using the reference matrix) to predict the mixture composition of probes. Binary classification is performed and finally, the predicted mixture composition is compared to the actual mixture composition. This process is repeated for each probe and the worst performing probe is removed until the overall classification is perfect. (B) Graphical representation of probe removal results. Individual probes are removed until the overall MCC value (confusion matrices shown on the right-hand side) is perfect (i.e equal to 1). (C) Workflow of sequential trimming of lists of good MuSIC probes. The final list of good MuSIC probes for single MuSIC probes (simulation 1) is used as the starting list for simulation 2. Then the final list of good MuSIC probes for barcodes (simulation 2) is used as the starting list for simulation 3.

We use three sequential sets of simulations to determine a list of “good” MuSIC probes that can be used (1) on their own, (2) for MuSIC barcodes (genetic screening), and (3) for two MuSIC barcodes (genetic interaction screening) (**Table S2**). The final list that is obtained in Simulation 1 is used for Simulation 2, and likewise 2 for 3 (**Fig. 3C**). For example, only probes that are good for use on their own are considered for MuSIC barcodes. The list of “good” MuSIC probes from Simulation 2 sets constraints on genetic screening for single gene effects and the list of good MuSIC probes from Simulation 3 sets constraints on genetic interaction screening.

The results of this process are summarized in **Table 1**. The final number of good MuSIC probes that can be unmixed with perfect classification for MuSIC barcodes and sets of two MuSIC barcodes are listed for each of the experimental setups (summarized in **Fig. 2B**). We found reasonable overlap between which probes were labeled as good between replicate runs (**Fig. S1**), although the overall number of probes seems to be a more reproducible and larger factor (**Table 1**). Given these results, the number of gRNA that can be used for genetic and genetic interaction screening are calculated from **Fig 1E**. In general, more lasers and lower noise allows for more probes and barcodes, as expected. For genetic screens, each scenario investigated suggested potential for genome-scale operation. For genetic interaction screens, 4 and 5 laser setups with low noise predicted operation at genome-scale. Even 2 and 3 laser setups with high noise predicted operation with 1000s of gRNA in genetic interaction screens, an order of magnitude above current methods. If we consider typical ranges of cell-to-cell variability in probe expression levels, then we can still generate gRNA on the same scale (**Table S3**). If we only consider MuSIC probes with one or two fluorescent proteins, as opposed to three, we can still achieve multiple hundred gRNA for genetic interaction screens (**Table S4**). Overall, these results suggest MuSIC barcoding theory represents a promising approach to transforming genetic perturbation technology.

**Table 1:**
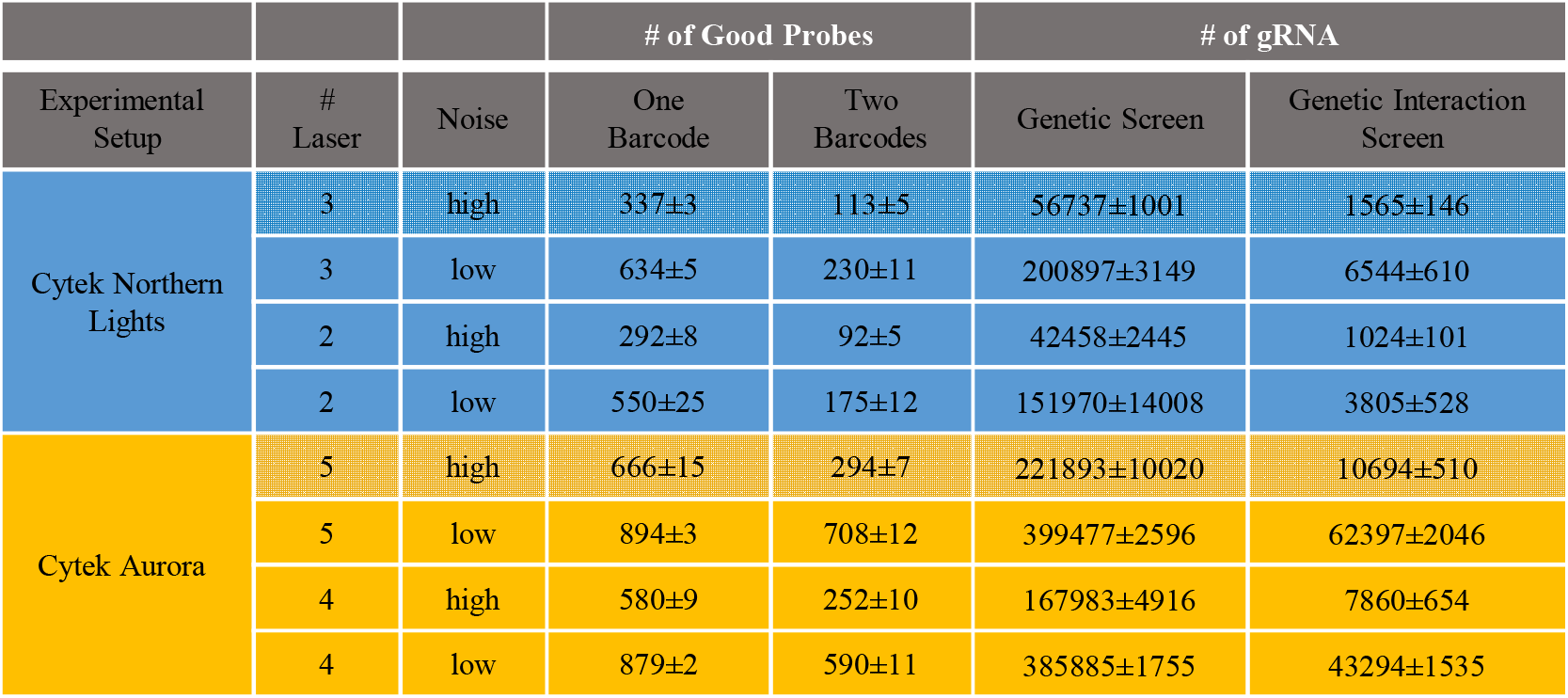
Simulated number of gRNA that could be used for genetic and genetic interaction screens. Results for the number of good probes that can be used to form barcodes and pairs of barcodes are shown for each experimental setup (the flow cytometer used), the number of lasers used, and the noise level (either low or high). Given the number of good probes, the number of potential gRNA for genetic and genetic interaction screens is listed. Results for the Cytek Northern Lights flow cytometer are highlighted in blue and results for the Aurora flow cytometer are highlighted in yellow.

|  |  |  | # of Good Probes |  | # of gRNA |  |
| --- | --- | --- | --- | --- | --- | --- |
| Experimental Setup | # Laser | Noise | One Barcode | Two Barcodes | Genetic Screen | Genetic Interaction Screen |
| Cytek Northern Lights | 3 | high | 337±3 | 113±5 | 56737±1001 | 1565±146 |
|  | 3 | low | 634±5 | 230±11 | 200897±3149 | 6544±610 |
|  | 2 | high | 292±8 | 92±5 | 42458±2445 | 1024±101 |
|  | 2 | low | 550±25 | 175±12 | 151970±14008 | 3805±528 |
| Cytek Aurora | 5 | high | 666±15 | 294±7 | 221893±10020 | 10694±510 |
|  | 5 | low | 894±3 | 708±12 | 399477±2596 | 62397±2046 |
|  | 4 | high | 580±9 | 252±10 | 167983±4916 | 7860±654 |
|  | 4 | low | 879±2 | 590±11 | 385885±1755 | 43294±1535 |

## Discussion

Here we propose an approach for single-cell, non-destructive, and potentially genome-scale genetic and genetic interaction screens. This work builds on our recently developed theory for Multiplexing using Spectral Imaging and Combinatorics (MuSIC). MuSIC probes are stably linked combinations of fluorophores with unique spectral signatures that can be deconvolved when in a mixture with other MuSIC probes. The novel concept introduced in this work is that of a MuSIC barcode, a combination of MuSIC probes. Given currently available fluorescent proteins, we estimate that ∼10^5^ unique MuSIC barcodes can be created from combinations of MuSIC probes. We devised a simulation workflow to generate lists of MuSIC probes that are likely to be deconvolvable in a mixture, given binary classification applications. These results show the potential for genetic screens at the human genome-scale and genetic interaction screens for at least 1000s of genes. In some cases (i.e. 4 or 5 lasers and low noise), results show the potential to perform genetic interaction screens at a human genome-scale.

What could be learned with non-destructive, single-cell genetic screens? When analyses are done on a single-cell level, each cell is analyzed independently, and as a result, multiple measurements can be done in parallel, increasing throughput^43–45^. To accomplish this, CRISPR screenings have been paired with single-cell RNA sequencing using methods like Perturb-Seq^46^, CRISP-seq^47^, or CROP-seq^48,49^. While single-cell sequencing has the ability to pair transcriptome responses to a nucleic acid barcode that indicates the genetic perturbation, it is as yet prohibitively expensive for covering interaction space^50,51^. Moreover, sequencing is a destructive technology so one cannot subsequently study perturbed cells-of-interest. The use of MuSIC barcodes could expand on the capabilities of these methods by allowing for high throughput genetic screening in a non-destructive manner. A non-destructive application in single live cells could allow sorting of rare cell types for subsequent follow-up studies. This could lead to co-isolating rare cell types thought to cooperate with each other for a disease phenotype.

What could be explored with high-dimensional non-destructive genetic interaction screens? One application is synthetic lethal interactions, which is defined as a genetic interaction that results in cell death, but disruption of the individual genes does not. Synthetic lethality has previously led to the discovery that poly(ADP-ribose) polymerase (PARP) inhibitors effectively kill BRCA1- and BRCA2 mutant tumor cells in breast cancer^52^. The proposed method may allow for genetic interaction screening at a near genome-scale, which could lead to the discovery of new synthetic lethal interactions in a high-throughput manner that is not currently possible. By discovering and exploiting synthetic lethal interactions in cancer cells, combinations of drugs can be used to treat cancer more effectively and at lower drug concentrations and thus lower toxicity^54^.

Although simulations suggest a large potential for the approach when applied to genetic screening, there are multiple technical hurdles to its implementation. How can one clone thousands of unique MuSIC barcodes specifically paired with matched gRNA? If one uses lentiviruses to deliver the constructs, how does one avoid template switching between genetically similar fluorescent proteins or barcodes, corrupting the connection between the barcode and gRNA^55^? The constructs may be large as well, so how does one achieve high enough titer to perform genetic interaction screening? Although flow cytometry is fast, can one assay enough cells to adequately explore gene interaction space? These are just some of the major issues that will arise, yet the potential applications, if these issues can be overcome, could be highly impactful.

Although we focused here on genetic screening as an application, genome-scale spectral barcode libraries could have other uses, such as high-dimensional cell lineage tracing. Current fluorescence-based lineage tracking is limited from spectral overlap and the number of unique probes. Techniques such as Brainbow work to fill this gap by using random ratios of different fluorophores to label cells^56^, but are still limited to ∼10s of deconvolvable colors^57^. This has been partially overcome through the use of DNA barcodes in each cell but requires destructive DNA sequencing to be deconvovled^57^. Music barcodes could be used to bridge this gap by expanding the available palette of color codes for fluorescence-based lineage tracing to potentially thousands of deconvolvable colors.

In conclusion, despite impending technical hurdles, the simulation studies presented here show the potential for MuSIC barcodes to enable high-dimensional genetic interaction screens at the human genome-scale. Its single-cell resolution compatibility and non-destructive features could also enable multiple new applications for established genetic screening, or for cell lineage tracking. The capabilities of this approach can further be increased by increasing the number of excitation lasers and/or the spectral wavelength resolution.

## Acknowledgments

The authors acknowledge funding from Clemson University, NIH/ NCI Grant R21CA196418, and NIH/NIGMS Grant R35 GM141891. M.E.M. received funding from the Department of Education Grant P200A180076. This work was performed using the Clemson University Palmetto Super Computing Cluster.

## Supplementary Figure and Table Legends

**Figure S1:**
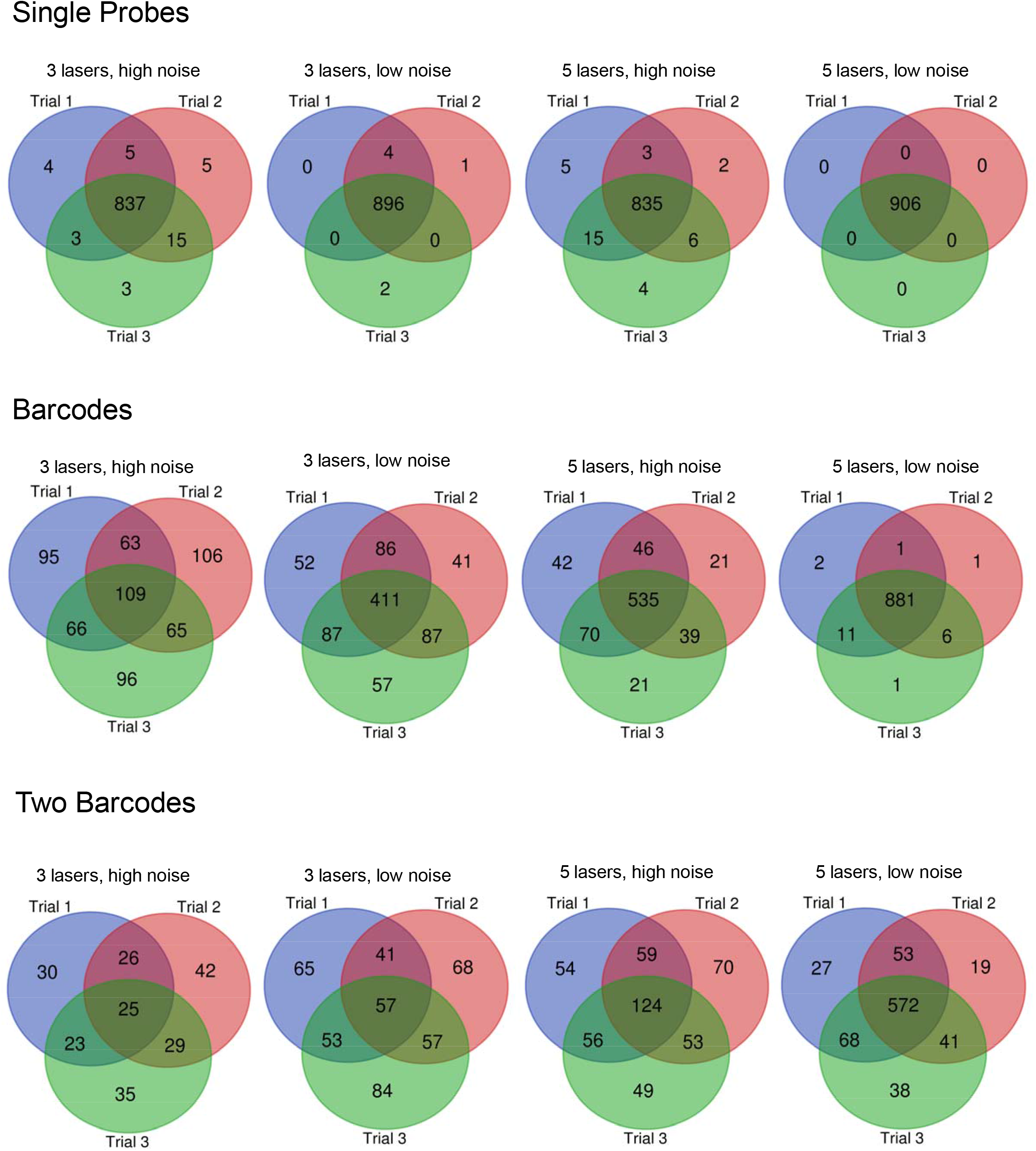
Comparison of lists of good probes between trials. The similarities and differences between the final lists of good probes for each trial are shown for each of the experimental setups.

Table S1: Fluorescent protein data. The maximum excitation and emission wavelength, brightness, extinction coefficient, and quantum yield for each fluorescent protein is found in the Attributes tab. Excitation and emission spectra for each fluorescent protein are found in the Excitation Spectra and Emission Spectra tabs, respectively. Sources for the raw data are found in the Sources tab.

Table S2: Probe lists. The lists of good probes for single probes, barcodes, and two barcodes are listed for each experimental setup in replicate.

**Table S3:** Simulated number of gRNA that could be used for genetic and genetic interaction screens with variable probe expression levels. We allowed probe expression levels to vary between 0.5 and 1.5 (relative) to capture single cell-to-cell variability. Results for the number of good probes that can be used to form barcodes and pairs of barcodes are shown for each experimental setup (the flow cytometer used), the number of lasers used, and the noise level (either low or high). Given the number of good probes, the number of potential gRNA for genetic and genetic interaction screens is listed.

|  |  |  | # of Good Probes |  | # of gRNA |  |
| --- | --- | --- | --- | --- | --- | --- |
| Experimental Setup | # Laser | Noise | One Barcode | Two Barcodes | Genetic Screen | Genetic Interaction Screen |
| Cytek Northern Lights | 3 | high | 339±16 | 119±7 | 57440±5403 | 1741±189 |
|  | 3 | low | 630±7 | 220±9 | 198184±4382 | 6014±474 |
|  | 2 | high | 264±5 | 87±4 | 34831±1371 | 922±91 |
|  | 2 | low | 538±16 | 164±6 | 144718±8708 | 3330±245 |
| Cytek Aurora | 5 | high | 619±15 | 299±2 | 191711±9555 | 11127±180 |
|  | 5 | low | 888±3 | 689±13 | 393539±2309 | 59156±2306 |
|  | 4 | high | 518±16 | 205±7 | 134331±8234 | 5198±367 |
|  | 4 | low | 870±6 | 537±10 | 377763±5355 | 35803±1334 |

**Table S4:** Simulated number of gRNA that could be used for genetic and genetic interaction screens when only considering one and two-way fluorescent protein probes. Probes containing three fluorescent proteins were not considered here. Results for the number of good probes that can be used to form barcodes and pairs of barcodes are shown for each experimental setup (the flow cytometer used), the number of lasers used, and the noise level (either low or high). Given the number of good probes, the number of potential gRNA for genetic and genetic interaction screens is listed.

|  |  |  | # of Good Probes |  | # of gRNA |  |
| --- | --- | --- | --- | --- | --- | --- |
| Experimental Setup | # Laser | Noise | One Barcode | Two Barcodes | Genetic Screen | Genetic Interaction Screen |
| Cytek Northern Lights | 3 | high | 100±3 | 61±5 | 1211±71 | 451±74 |
|  | 3 | low | 113±1 | 90±4 | 1559±10 | 979±80 |
|  | 2 | high | 88±1 | 50±3 | 932±38 | 302±38 |
|  | 2 | low | 109±2 | 69±2 | 1450±48 | 573±22 |
| Cytek Aurora | 5 | high | 104±2 | 91±2 | 1310±45 | 1006±53 |
|  | 5 | low | 113±0 | 109±1 | 1559±19 | 1449±36 |
|  | 4 | high | 102±2 | 81±2 | 1259±34 | 781±39 |
|  | 4 | low | 114±0 | 109±1 | 1596±0 | 1431±31 |

