## Supplementary material for "A Theory for High-Throughput Genetic Interaction Screening": Code: BinClass.m

```
function fhatbin = BinClass(fhatmat,Thresh)
%inputs:    Thresh- vector containing different thresholds for determining
%           whether or not a probe is present
%           fhatmat- Matrix containing fhat vectors for each probe. dim: (probes, probes*n) 
%outputs:   fhatbin- 3D matrix containing binary fhat vector for each probe
%           at each threshold. dim( probes, fhatbin, thresh)
%purpose:   generate fhat bin for particular noise level and laser. test each
%           fhatmat for each threshold configuration
parfor i = 1:length(Thresh)
    fhatbin(:,:,i) = fhatmat > Thresh(i); % set value equal to 1 if value in fhatmat is greater than the current threshold 
end

    
```
