## Supplementary material for "A Theory for High-Throughput Genetic Interaction Screening": Code: calc_confusion.m

```
function conf_mats = calc_confusion(fhatbin,fmat)
%inputs:    fhatbin- 3D matrix containing binary fhat vector for each probe
%           at each threshold. dim:(probes, fhatmat, thresh)
%           fmat- real f matrix for each probe 
%outputs:   conf_mats- confusion matrix 
%purpose:   generate confusion matrix for each probe at each threshold 
N_Probes = length(fhatbin(:,1,1));
N_Thresh = length(fhatbin(1,1,:));
fmat=logical(fmat);
parfor i=1:N_Probes
    for j=1:N_Thresh
        conf_mats{i,j}=confusionmat(fmat(i,:),fhatbin(i,:,j));
    end
end

% need to generate conf mats for each of the 4 fhatbins 
```
