## Supplementary material for "A Theory for High-Throughput Genetic Interaction Screening": Code: Calc_R0.m

```
function [R0] = Calc_R0(Ex, Em, Q_D, e_A,lam)
%Inputs: excitation spectra, mission spectra, donor quantum yield, acceptor
%extinction coefficient, wavelengths
%Outputs: Forster Radius
K2 = 2./3; %Kappa squared 
beta = 6.33e-6; %(M*nm^3)
J=trapz((lam).^4.*Em.*Ex); %(nm^4) overlap integral
R0=(beta .* K2 .* Q_D .* e_A .*J).^(1/6); %(nm) Forster radius
```
