## Supplementary material for "A Theory for High-Throughput Genetic Interaction Screening": Code: CalcIntensity1F.m

```
function I=CalcIntensity1F(C,B,Ex,Em,lamv,exlam,Iex)

f=Ex(lamv==exlam); %determine which parts of excitation spectra overlap with excitation light
I=f.*C.*B.*Em.*Iex; %emission intensity proportional to excitation strength (f), probe concentration (C), brightness (B), emission intensity (Em), and intensity of the excitation light (normally 1 here)
```
