## Supplementary material for "A Theory for High-Throughput Genetic Interaction Screening": Code: CalcIntensity2F_2.m

```
function I=CalcIntensity2F_2(C,B1,B2,Ex1,Ex2,Em1,Em2,lamv,exlam,Iex)

E=0.5;
f1=Ex1(lamv==exlam); %determine which parts of excitation spectra overlap with excitation light
f2=Ex2(lamv==exlam); %determine which parts of excitation spectra overlap with excitation light
Eadj=E*(1-f2); %reduce FRET efficiency in places where there is direct acceptor excitation

I1D=Iex.*f1.*(1-Eadj).*C.*B1.*Em1;  %intensity of the donor emission (subtracting energy lost to FRET)
I1A=Iex.*f1.*Eadj.*C.*B2.*Em2;  %intensity of the acceptor emission due to FRET
I1Ad=Iex.*f2.*(1-Eadj).*C.*B2.*Em2; %intensity of the acceptor emission due to direct excitation, correcting for fraction of molecules "FRETing"

I=I1D+I1A+I1Ad; %add 'em all up
```
