## Supplementary material for "A Theory for High-Throughput Genetic Interaction Screening": Code: CalcIntensity3F_2.m

```
function I=CalcIntensity3F_2(C,B1,B2,B3,Ex1,Ex2,Ex3,Em1,Em2,Em3,lamv,exlam,Iex)

E1=.5;
E2=.5;
%determine which parts of excitation spectra overlap with excitation light
f1=Ex1(lamv==exlam); 
f2=Ex2(lamv==exlam);
f3=Ex3(lamv==exlam);
%reduce FRET efficiency in places where there is direct acceptor excitation
Eadj1=E1*(1-f2);
Eadj2=E2*(1-f3);


I1D=Iex.*f1.*(1-Eadj1).*C.*B1.*Em1; %intensity of the first donor emission (subtracting energy lost to FRET)
I1A1=Iex.*f1.*Eadj1.*C.*B2.*Em2; %intensity of the first acceptor emission due to FRET from first donor
I1A2=Iex.*f1.*Eadj1.*Eadj2.*C.*B3.*Em3; %intensity of the second acceptor emission due to FRET from first donor
I1D2=Iex.*f2.*(1-Eadj1-Eadj2).*C.*B2.*Em2; %intensity of the first acceptor emission due to direct excitation, correcting for fraction of molecules "FRETing"
I1A2_2=Iex.*f2.*Eadj2.*C.*B3.*Em3; %intensity of the second acceptor emission due to FRET from the second donor
I1A2d=Iex.*f3.*(1-Eadj2-Eadj1.*Eadj2).*C.*B3.*Em3; %intensity of the second acceptor emission due direct excitation, correction for fraction of molecules "FRETing"

I=I1D+I1A1+I1A2+I1D2+I1A2_2+I1A2d;
```
