## Supplementary material for "A Theory for High-Throughput Genetic Interaction Screening": Code: CalcOverallClassificationMetrics.m

```
function [MCC_Overall, Precision, Recall, Percent_Correct]=CalcOverallClassificationMetrics(fmat,fhatmat,threshs)
%Inputs:    fmat- the actual levels. dim: (probes, # experiments)
%           fhatmat- estimated levels. dim: (probes, # experiments)
%           threshs- binary classification threshold. dim: (1,probes)
%Output:    MCC_Overall- calculates MCC based on aggregation over all probes and
%           experiments
%           Precision- Calculates precision over all probes and experiments
%           Recall- Calculates recall over all probes and experiments
%           Percent_Correct- Calculates the precent correct over all probes and experiments
%Purpose:   Generate classification metrics for a group of probes

%Perform Binary Classification
for i=1:length(fmat(1,:))
    fhatbin(:,i) = fhatmat(:,i) > threshs';
end

%Vectorize
fhatbinv=reshape(fhatbin,[],1);
fmatv=logical(reshape(fmat,[],1));

%Calculate metrics
mat=confusionmat(fmatv,fhatbinv)
MCC_Overall = (mat(2,2)*mat(1,1)-mat(1,2)*mat(2,1))/sqrt((mat(2,2)+mat(1,2))*(mat(2,2)+mat(2,1))*(mat(1,1)+mat(1,2))*mat(1,1)+mat(2,1))
% << for an existing Confision Matrix 'ConfusionMat' >>
Precision = mat(2,2)/(mat(2,2)+mat(1,2))
% And another for recall
Recall = mat(2,2)/(mat(2,2)+mat(2,1))
Percent_Correct=(mat(1,1)+mat(2,2))/(mat(2,2)+mat(1,1)+mat(1,2)+mat(2,1))
```
