## Supplementary material for "A Theory for High-Throughput Genetic Interaction Screening": Code: condensed_refs.m

```
function R_New = condensed_refs(Exlam, R, Bin,lam)
%inputs:    Exlam - excitation wavelengths
%           R - reference matrix at every nm 
%           Bin - binning structure for flow cytometer 
%           lam - corresponding wavelengths for reference matrix 
%outputs:   R_New - condensed refernece matrix 
%Purpose:   creates a condensed reference matrix according to flow cytometer
%           binning 
R_New = zeros(length(Bin(:,1)),length(R(1,:)));
for i = 1:length(R(1,:)) %0 to # columns in R --> number of probes in ref matrix 
    R_New(:,i) = observation(Exlam,R(:,i),Bin,lam);
end
    
```
