## Supplementary material for "A Theory for High-Throughput Genetic Interaction Screening": Code: CreateAllPossibleProbes.m

```
function [probes]=CreateAllPossibleProbes(Ex,Em,lam,QY,ext,R0_Thresh)
%Inputs:    Ex- individual FP excitation spectra
%           Em- individual FP emission spectra
%           lam- wavelengths
%           QY- quantum yield
%           ext- extinction coefficient
%           R0_Thresh- cut-off for minimum R0
%Output:    probes- matrix where rows are individual probes, columns are FP
%           monomers, and an element of 1 means that FP is in that probe
%Purpose:   Generate list of probes above a specified FRET efficiency 

N=length(Ex(1,:));

Counter=1;
for i=1:N
    %single probes
    probes(Counter,i)=1;
    Counter=Counter+1;
    for j=1:N
        if Calc_R0(Ex(:,j),Em(:,i),QY(i),ext(j),lam)>R0_Thresh
        %double probes
            probes(Counter,i)=1;
            probes(Counter,j)=1;
            Counter=Counter+1;
%         for k=1:N %only if we find a suitable double, consider a triple
%             if Calc_R0(Ex(:,j),Em(:,i),QY(i),ext(j),lam)>R0_Thresh
%                 %triple probes
%                 probes(Counter,i)=1;
%                 probes(Counter,j)=1;
%                 probes(Counter,k)=1;
%                 Counter=Counter+1;
%             end
%         end
        end
    end
end
%remove any duplicates (not worrying about order in protein for now)
[probes,~,~] = unique(probes, 'rows');
   
```
