## Supplementary material for "A Theory for High-Throughput Genetic Interaction Screening": Code: f_and_fhat.m

```
function [fhatmat,fmat]=f_and_fhat(probes,R,noise,ntrials)
%inputs: probes - fluorescent proteins combinations that make up each probe 
%        R - emission spectra of each probe for each excitation channel 
%        noise - standard deviation of normal distribution (zero mean)
%        ntrials - number of replicates for each probe
%output: fhatmat - predicted propbes present in mixture
%        fmat - actual probes present in mixture
%Purpose: generate f and fhat data for each probe 
N_PROBES=length(probes(:,1));
n=ntrials;
fhatmat = zeros(N_PROBES,N_PROBES*n);
fmat = zeros(N_PROBES,N_PROBES*n);
    for j=1:N_PROBES
        for i = 1:n
        f = zeros(N_PROBES,1);
        f(j)=1;
        mu = R * f ; %calculate mu from ref matrix and f
        Length_mu = length(mu); 
        rng('shuffle');
        temp=noise.*randn(Length_mu,1); %creates variable for noise
        mu = mu + temp; %add noise to mu
        f_hat = Unmix(R,mu,0,zeros(N_PROBES,1),[],1);
        fmat(:,i+n*(j-1)) = f;
        fhatmat(:,i+n*(j-1)) = f_hat;
        f_hat = zeros(N_PROBES,1);
        end
    end
```
