## Supplementary material for "A Theory for High-Throughput Genetic Interaction Screening": Code: f_and_fhat_barcodes_hybrid.m

```
function [fhatmat,fmat]=f_and_fhat_barcodes_hybrid(probes,R,noise,ntrials)
%inputs: probes - fluorescent proteins combinations that make up each probe 
%        R - emission spectra of each probe for each excitation channel 
%        noise - standard deviation of normal distribution (zero mean)
%        ntrials - number of replicates for each barcode
%output: fhatmat - predicted propbes present in mixture
%        fmat - actual probes present in mixture
%Purpose: generate f and fhat data for each barcode

N_PROBES=length(probes(:,1));
n=ntrials;
for i = 1:n
    parfor j = 1:N_PROBES
        rng('shuffle');
        f = zeros(N_PROBES,1);
        f(j)=1;
        f(randi([1 N_PROBES]))=1;
        mu = R * f ; %calculate mu from ref matrix and f
        Length_mu = length(mu); 
        temp=noise.*randn(Length_mu,1); %creates variable for noise
        mu = mu + temp; %add noise to mu
        f_hat = Unmix(R,mu,0,zeros(N_PROBES,1),[],1);
        fmat(:,j,i) = f;
        fhatmat(:,j,i) = f_hat;
        f_hat = zeros(N_PROBES,1);
    end
end
fmat=reshape(fmat,N_PROBES,N_PROBES*n);
fhatmat=reshape(fhatmat,N_PROBES,N_PROBES*n);
```
