## Supplementary material for "A Theory for High-Throughput Genetic Interaction Screening": Code: GenerateProbeData_2L_HN.m

```
% Generate good probes for 2 laser-high noise
clear
load SingleProbeVariables.mat %all probes
%load SingleProbeVariables_short.mat %only 1 & 2-way probes
% Single Probes
[OverallMCC_probes_HN_2l,ProbesRemoved_probes_HN_2l,ProbesRemoved_names_probes_HN_2l,Opt_Thresh_probes_HN_2l,Good_probes_probes_HN_2l,R_Cond_final_probes_HN_2l] = RemoveProbes_onebyone(probes,R_2l,high_noise,thresh,names);
% Barcode
[OverallMCC_barcodes_HN_2l,ProbesRemoved_barcodes_HN_2l,Opt_Thresh_probes_barcodes_HN_2l,Good_probes_final_barcodes_HN_2l,R_Cond_barcodes_HN_2l_final,Probes_Removed_names_barcodes_HN_2l] = RemoveBarcodes_onebyone(Good_probes_probes_HN_2l,R_Cond_final_probes_HN_2l,high_noise,thresh,names);
% Two Barcodes
[OverallMCC_twobarcodes_HN_2l,ProbesRemoved_twobarcodes_HN_2l,Opt_Thresh_probes_twobarcodes_HN_2l,Good_probes_final_twobarcodes_HN_2l,R_Cond_Barcodes_final_twobarcodes_HN_2l,Probes_Removed_names_twobarcodes_HN_2l] = RemoveTwoBarcodes_onebyone(Good_probes_final_barcodes_HN_2l,R_Cond_barcodes_HN_2l_final,high_noise,thresh,names)
save('ProbeData_2l_HN')
```
