## Supplementary material for "A Theory for High-Throughput Genetic Interaction Screening": Code: GenerateProbeData_4L_LN.m

```
% Generate good probes for 4 laser-low noise
clear
load SingleProbeVariables.mat %all probes
%load SingleProbeVariables_short.mat %only 1 & 2-way probes
% Single Probes
[OverallMCC_probes_LN_4l,ProbesRemoved_probes_LN_4l,ProbesRemoved_names_probes_LN_4l,Opt_Thresh_probes_LN_4l,Good_probes_probes_LN_4l,R_Cond_final_probes_LN_4l] = RemoveProbes_onebyone(probes,R_4l,low_noise,thresh,names);
% Barcodes
[OverallMCC_barcodes_LN_4l,ProbesRemoved_barcodes_LN_4l,Opt_Thresh_probes_barcodes_LN_4l,Good_probes_final_barcodes_LN_4l,R_Cond_barcodes_LN_4l_final,Probes_Removed_names_barcodes_LN_4l] = RemoveBarcodes_onebyone(Good_probes_probes_LN_4l,R_Cond_final_probes_LN_4l,low_noise,thresh,names);
% Two Barcodes
[OverallMCC_twobarcodes_lownoise_4l,ProbesRemoved_twobarcodes_lownoise_4l,Opt_Thresh_probes_twobarcodes_lownoise_4l,Good_probes_final_twobarcodes_lownoise_4l,R_Cond_Barcodes_final_twobarcodes_lownoise_4l,Probes_Removed_names_twobarcodes_LN_4l] = RemoveTwoBarcodes_onebyone(Good_probes_final_barcodes_LN_4l,R_Cond_barcodes_LN_4l_final,low_noise,thresh,names)
save('ProbeData_4l_LN')
```
