## Supplementary material for "A Theory for High-Throughput Genetic Interaction Screening": Code: GenerateProbeData_5L_HN.m

```
% Generate good probes for 5 laser-high noise
clear
load SingleProbeVariables.mat %all probes
%load SingleProbeVariables_short.mat %only 1 & 2-way probes
% Single Probes
[OverallMCC_probes_HN_5l,ProbesRemoved_probes_HN_5l,ProbesRemoved_names_probes_HN_5l,Opt_Thresh_probes_HN_5l,Good_probes_probes_HN_5l,R_Cond_final_probes_HN_5l] = RemoveProbes_onebyone(probes,R_5l,high_noise,thresh,names);
% Barcodes
[OverallMCC_barcodes_HN_5l,ProbesRemoved_barcodes_HN_5l,Opt_Thresh_barcodes_HN_5l,Good_probes_barcodes_HN_5l,R_Cond_final_barcodes_HN_5l,ProbesRemoved_names_barcodes_HN_5l] = RemoveBarcodes_onebyone(Good_probes_probes_HN_5l,R_Cond_final_probes_HN_5l,high_noise,thresh,names);
% Two Barcodes
[OverallMCC_twobarcodes_HN_5l,ProbesRemoved_twobarcodes_HN_5l,Opt_Thresh_twobarcodes_HN_5l,Good_probes_twobarcodes_HN_5l,R_Cond_final_twobarcodes_HN_5l,ProbesRemoved_names_twobarcodes_HN_5l] = RemoveTwoBarcodes_onebyone(Good_probes_barcodes_HN_5l,R_Cond_final_barcodes_HN_5l,high_noise,thresh,names);
save('ProbeData_5l_HN')
```
