## Supplementary material for "A Theory for High-Throughput Genetic Interaction Screening": Code: Get_OptThresh.m

```
function [Opt_Thresh,MCCMax] = Get_OptThresh(MCC_Mat,Thresh)
%Inputs:    MCC_Mat- matrix of MCC values for each probe
%           Thresh- vector of thresholds for binary classification 
%Outputs:   Opt_Thresh- optimum threshold for binary classification for each
%           probe. dim: (probes)
%           MCCMax- vector containing the maximum MCC value for each probe.
%           dim (probes)
%Purpose:   determine the optimum threshold for binary classification for
%           each probe
[MCCMax,MCCIn] = max(MCC_Mat'); %MCCMax returns horiztontal vector of max MCC value for each probe
                                %MCCIn returns which column the max MCC value was in 
Opt_Thresh = Thresh(MCCIn); %determines conresponding threshold
```
