## Supplementary material for "A Theory for High-Throughput Genetic Interaction Screening": Code: GetEmSubVector.m

```
function EmSubVector=GetEmSubVector(Em_Sim, C_Ex, N_Ex)
%Inputs:    Em_Sim - simulated emission spectra at every nm
%           C_Ex  -  current excitation channel
%           N_Ex - number of excitation channels
%Outputs:   EmSubVector - portion of emission spectra for particular excitation channel 
WL_Range = length(Em_Sim(:,1))/N_Ex; %must have same number of wavelengths for each excitation channel
EmSubVector=Em_Sim(1+WL_Range*(C_Ex-1):WL_Range*C_Ex,1);
```
