## Supplementary material for "A Theory for High-Throughput Genetic Interaction Screening": Code: GetIndividualProbeNames.m

```
function [probenames]=GetIndividualProbeNames(names,probes)

%creates a list of probe names

for i=1:length(probes(:,1))
    
    ind=find(probes(i,:)==1);
    name='';
    for j=1:length(ind)
        if j==length(ind)
            name=strcat(name,names(ind(j)));
        else
            name=strcat(name,names(ind(j)),'-');
        end
    end
    probenames{i}=name;
end
probenames=probenames';
```
