## Supplementary material for "A Theory for High-Throughput Genetic Interaction Screening": Code: GetMCC.m

```
function MCC_Mat = GetMCC(conf_mats)
%input :    matrix of confusion matrix for each probe at each threshold 
%Output:    matrix of MCC values for each confusion matrix. dim: (conf_mats
MCC_Mat = zeros(size(conf_mats));
for i = 1 : length(conf_mats(:,1))
    for j = 1 : length(conf_mats(1,:))
        mat = conf_mats{i,j};
        MCC_Mat(i,j) = (mat(2,2)*mat(1,1)-mat(1,2)*mat(2,1))/sqrt((mat(2,2)+mat(1,2))*(mat(2,2)+mat(2,1))*(mat(1,1)+mat(1,2))*mat(1,1)+mat(2,1));
    end
end
```
