## Supplementary material for "A Theory for High-Throughput Genetic Interaction Screening": Code: GetRefMat.m

```
function R=GetRefMat(Em,Ex,lam,B,probes,Exlam)

N_PROBES=length(probes(:,1));
N_EX=length(Exlam);
Iex=ones(N_EX,1);  %relative intensity of excitation at certain wavelengths, typically not considered here and taken as 1
N_LAM=length(Em(:,1));
R=zeros(N_LAM*N_EX,N_PROBES);
for i=1:N_EX
    for j=1:N_PROBES
        ind=find(probes(j,:)==1);
        switch sum(probes(j,:))
            case 1  %single probes
                R((i-1)*N_LAM+1:i*N_LAM,j)=CalcIntensity1F(1,B(ind),Ex(:,ind),Em(:,ind),lam,Exlam(i),Iex(i));
            case 2  %double probes
                R((i-1)*N_LAM+1:i*N_LAM,j)=CalcIntensity2F_2(1,B(ind(1)),B(ind(2)),Ex(:,ind(1)),Ex(:,ind(2)),Em(:,ind(1)),Em(:,ind(2)),lam,Exlam(i),Iex(i));
            case 3  %triple probes
                R((i-1)*N_LAM+1:i*N_LAM,j)=CalcIntensity3F_2(1,B(ind(1)),B(ind(2)),B(ind(3)),Ex(:,ind(1)),Ex(:,ind(2)),Ex(:,ind(3)),Em(:,ind(1)),Em(:,ind(2)),Em(:,ind(3)),lam,Exlam(i),Iex(i));
        end
    end
end
```
