## Supplementary material for "A Theory for High-Throughput Genetic Interaction Screening": Code: observation.m

```
function Em_New=observation(Exlam, Em_Sim, Bin,lam)
%Inputs: Exlam - excitation wavelengths
%   Em_Sim - simulated emission spectra 
%   Bin - binning matrix with ex channel, emission start wl, emission end
%   lam - wavelengths
%Outputs: Em_New - condensed emission spectra
%Purpose: creates a condensed emission spectra according to flow cyte binning 
N_Ex=length(Exlam); % number of excitation channels 
Em_New = zeros(length(Bin(:,1)),1);
for i=1:length(Bin(:,1)) % cycle through rows in bin
 Em_SimTemp=GetEmSubVector(Em_Sim,Bin(i,1),N_Ex);% get portion on Em_Sim for particular Excitation channel   
 inds=find(lam>=Bin(i,2)&lam<=Bin(i,3));%could use logical indexing
 Em_New(i) = mean(Em_SimTemp(inds));
end 

        
    
```
