## Supplementary material for "A Theory for High-Throughput Genetic Interaction Screening": Code: READ_ME.txt

A Theory for High-Throughput Genetic Interaction Screening 

Madeline E. McCarthy, William B. Dodd, Xiaoming Lu, Nishi Patel, Charlotte Haskell, Hugo Sanabria, Mark A. Blenner, and Marc R. Birtwistle

1.) Inputs for probe simulations are found in SingleProbeVariables.mat. Inputs for probe simulations for only one- and two-way probes are found in SingleProbeVariables_short.mat.

2.) These inputs can be loaded into the scripts GenerateProbeData_3l_HN.m, GenerateProbeData_3l_LN.m, GenerateProbeData_5l_HN.m, and GenerateProbeData_5l_LN.m 

3.) The outputs for these scripts contain the lists of good probes for single probes, barcodes, and two barcodes along with the lists of probes removed in each round. 
