## Supplementary material for "A Theory for High-Throughput Genetic Interaction Screening": Code: RemoveBarcodes_onebyone.m

```
function [OverallMCC,ProbesRemoved,Opt_Thresh_probes,Good_probes_final,R_Cond_Barcodes_final,ProbesRemoved_names] = RemoveBarcodes_onebyone(Good_probes,R_Cond_final,noise,thresh,names)
%Inputs:    Good_probes- List of good probes from RemoveProbes_onebyone
%           R_Cond_final- Condensed reference matrix for Good_probes
%           noise- noise level 
%           thresh- vector of thresholds for binary classification
%           names- list of names for fluorescent proteins 
%Outputs:   OverallMCC- Vector containing the overall MCC value from each loop
%           ProbesRemoved- List of probes removed in each loop
%           Opt_Thresh_probes- Vector containing the optimum threshold for binary classification for each probe
%           Good_probes_final- list of final good probes (Overall MCC = 1)
%           R_Cond_Barcodes_final- Condensed reference matrix for Good_probes_final
%           ProbesRemoved_names- list of probe names for Good_probes_final
ProbesRemoved = zeros(size(Good_probes));
OverallMCC = zeros(length(Good_probes),1);
    for i=1:length(Good_probes)
        [fhatmat_barcodes,fmat_barcodes]=f_and_fhat_barcodes_hybrid(Good_probes,R_Cond_final,noise,3);
 %       [fhatmat_barcodes,fmat_barcodes]=f_and_fhat_barcodes_hybrid_rnd_conc(Good_probes,R_Cond_final,noise,3);
 %          use for variable probe expression levels
        fhatbin_barcodes = BinClass(fhatmat_barcodes,thresh);
        conf_mats_barcodes = calc_confusion(fhatbin_barcodes,fmat_barcodes);
        MCC_Mat_barcodes = GetMCC(conf_mats_barcodes);
        [Opt_Thresh_probes,MCCMax_barcodes] = Get_OptThresh(MCC_Mat_barcodes,thresh);
        [MCC_Overall]=CalcOverallClassificationMetrics(fmat_barcodes,fhatmat_barcodes,Opt_Thresh_probes);
        [~,MinInd] = min(MCCMax_barcodes);
        ProbesRemoved(i,:) = Good_probes(MinInd,:);
        Good_probes(MinInd,:)=[];
        OverallMCC(i)=(MCC_Overall);
        R_Cond_final(:,MinInd)=[];
        if MCC_Overall == 1
            break
        end 
    end
    ProbesRemoved_names=GetIndividualProbeNames(names,ProbesRemoved);
    [fhatmat_barcodes,fmat_barcodes]=f_and_fhat_barcodes_hybrid(Good_probes,R_Cond_final,noise,3);
    fhatbin_barcodes = BinClass(fhatmat_barcodes,thresh);
    conf_mats_barcodes = calc_confusion(fhatbin_barcodes,fmat_barcodes);
    MCC_Mat_barcodes = GetMCC(conf_mats_barcodes);
    [Opt_Thresh_probes] = Get_OptThresh(MCC_Mat_barcodes,thresh);
    R_Cond_Barcodes_final = R_Cond_final;
    Good_probes_final=Good_probes;
```
