## Supplementary material for "A Theory for High-Throughput Genetic Interaction Screening": Code: RemoveProbes_onebyone.m

```
function [OverallMCC,ProbesRemoved,ProbesRemoved_names,Opt_Thresh,Good_probes,R_Cond_final] = RemoveProbes_onebyone(probes,R_Cond,noise,thresh,names)
%Inputs:    probes- List probes
%           R_Cond- Condensed reference matrix for probes
%           noise- noise level 
%           thresh- vector of thresholds for binary classification
%           names- list of names for fluorescent proteins 
%Outputs:   OverallMCC- Vector containing the overall MCC value from each loop
%           ProbesRemoved- List of probes removed in each loop
%           ProbesRemoved_names- list of probe names for Good_probes
%           Opt_Thresh- Vector containing the optimum threshold for binary classification for each probe
%           Good_probes- list of final good probes (Overall MCC = 1)
%           R_Cond_final- Condensed reference matrix for Good_probes_final
%           
ProbesRemoved = zeros(size(probes));
OverallMCC = zeros(length(probes),1);
    for i=1:length(probes)
        [fhatmat,fmat]=f_and_fhat(probes,R_Cond,noise,10);
        %[fhatmat,fmat]=f_and_fhat_rnd_conc(probes,R_Cond,noise,10);
            % use for variable probe expression levels
        fhatbin = BinClass(fhatmat,thresh);
        conf_mats = calc_confusion(fhatbin,fmat);
        MCC_Mat = GetMCC(conf_mats);
        [Opt_Thresh,MCCMax] = Get_OptThresh(MCC_Mat,thresh);
        [MCC_Overall]=CalcOverallClassificationMetrics(fmat,fhatmat,Opt_Thresh);
        [~,MinInd] = min(MCCMax);
        ProbesRemoved(i,:) = probes(MinInd,:);
        probes(MinInd,:)=[];
        OverallMCC(i)=(MCC_Overall);
        R_Cond(:,MinInd)=[];
        if MCC_Overall == 1
            break
        end
    end
    ProbesRemoved_names=GetIndividualProbeNames(names,ProbesRemoved);
    [fhatmat,fmat]=f_and_fhat(probes,R_Cond,noise,1);
    fhatbin = BinClass(fhatmat,thresh);
    conf_mats = calc_confusion(fhatbin,fmat);
    MCC_Mat = GetMCC(conf_mats);
    [Opt_Thresh] = Get_OptThresh(MCC_Mat,thresh);
    R_Cond_final = R_Cond;
    Good_probes = probes;
    

    
```
