## Supplementary material for "A Theory for High-Throughput Genetic Interaction Screening": Code: Reshapeafterparallel.m

```
function fnew=Reshapeafterparallel(f,n,N_PROBES)

%puts the parallel generated cell array version of barcode simulation data
%into the "old" format of probes by experiments

Counter=1;
for k=1:N_PROBES
    for j=k:N_PROBES
        for i = 1:n
            fnew(:,Counter)=f(i,j,k);
            Counter = Counter + 1;
        end
    end
end
```
