## Supplementary material for "A Theory for High-Throughput Genetic Interaction Screening": Code: Unmix.m

```
function [Levels]=Unmix(Refs,Samples,Sum2One,LB,UB,RefSums)

%Make Sum2One = 0 if you don't want it, or = 1 if you do; this refers to a
%constraint where the estimated levels of all probes should sum to 1

%Set LB = [] and UB = [] if you don't want any

Aeq=[];beq=[];
if Sum2One
    Aeq=ones(1,length(Refs(1,:))); beq=1;
end
if isempty(RefSums)
    RefSums=ones(1,length(Refs(1,:)));
end
for i=1:length(Samples(1,:))
    options = optimoptions('lsqlin','Display','none');
    Levels(:,i)=lsqlin(Refs,Samples(:,i),[],[],Aeq,beq,LB,UB,[],options)./RefSums';
    %Levels(:,i)=Levels(:,i)./sum(Levels(:,i));
end
```
